# Seeing halos: Spatial and consumer-resource constraints to landscape of fear patterns

**DOI:** 10.1101/2024.04.15.587800

**Authors:** Theresa W. Ong, Lisa C. McManus, Vítor V. Vasconcelos, Luojun Yang, Chenyang Su

**Author notes:** Co-first authors (Theresa W. Ong, Lisa C. McManus, Vítor V. Vasconcelos).

## Abstract

Coral halos are algae-free space surrounding coral patches linked to the landscape of fear hypothesis, where risk-averse herbivores avoid grazing too far from coral shelter. Halo patterns thus provide insight into the state of overfished predator populations on reefs. However, halos are absent in some protected reefs and their sizes are uncorrelated with predator abundance. Here, we show that these observations are consistent with the landscape of fear hypothesis. When coral are dispersed, herbivores have more even shelter from predators across the seascape, allowing herbivores to overgraze and halos to oscillate. When coral are clustered, shelter is limited and halos can stabilize, but reducing herbivore limitation drives critical transitions to cycles that are difficult to recover from. Discordance between theory and observations may, therefore, arise from differences in underlying coral spatial distributions, with broad implications for how the landscape of fear influences pattern formation and ecological resilience in patchy ecosystems.

## Introduction

Coral reefs can exhibit a distinctive spatial feature known as grazing or sand halos, characterized by coral patches that are separated from surrounding algae or seagrass by a “conspicuous band of bare sand” in the form of a ring (Randall, 1965). These halos have been documented since the 1960s and are visible from satellite images (Ogden et al., 1973) (Fig. 1). Recent studies suggest that coral halos may signal the presence of a robust reef community composed of healthy populations of herbivores and higher trophic levels that may also restrict bioturbation by prey and enhance carbon sequestration on reefs (Atwood et al., 2018; Madin et al., 2019b). Coral reefs are experiencing unprecedented global declines due to a confluence of global change factors including the extirpation of many top predators from our oceans (Sully et al., 2022). Thus, using coral halos for remote assays of reef community and ecosystem health holds significant conservation potential. However, the underlying biological and physical mechanisms that produce these halos remain unresolved, especially interactions between coral spatial clustering and consumer-resource dynamics.

**Figure 1:**
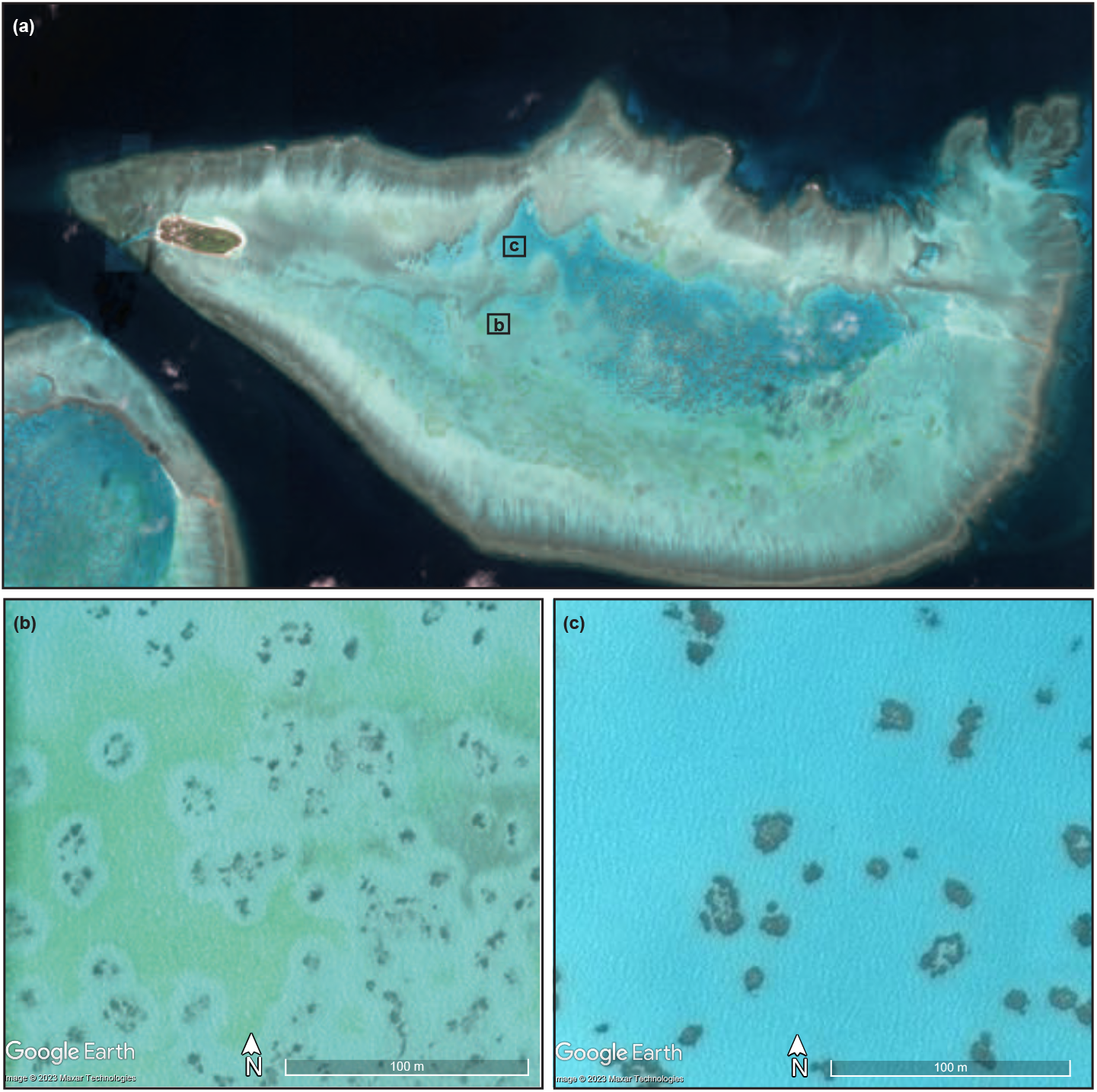
Coral halos are visible in satellite imagery of (a) Heron Island on the Great Barrier Reef, Australia. Halos are bands of algae-free space surrounding coral heads due to grazing. Close up view of selected reef regions indicated by labeled squares in (a), which are drawn at scale of panels (b) where there are clear halos and (c) no halos. Satellite imagery dated 2016 is from Maxar Technologies downloaded using Google Earth in 2023.

Large-scale spatial ecological patterns like coral halos often emerge from endogenous interactions among individuals, frequently involving multiple species. These patterns can reflect changes in the underlying conditions of an ecosystem and can serve as early warning indicators of shifts in ecosystem health and resilience (Scheffer, 2009). Ecological resilience refers to the ability of an ecosystem to sustain, recover, and adapt to perturbations within a specific ecological context (Holling, 1973). In the Namib Desert, for example, regularly dispersed bare spots surrounded by grass rings are likely generated through a combination of distance-dependent feedbacks within the plant community and competition among termite colonies (Bonachela et al., 2015; Tarnita et al., 2017). These and other vegetation patterns may serve as early warning indicators of a catastrophic shift from high plant abundance to a barren state as rainfall decreases (Rietkerk et al., 2004). However, the qualitative nature of this response is context-dependent. In the Namib Desert, systems with termites are more resilient to drought because plants can persist and recover under reduced rainfall (Bonachela et al., 2015). In coral reefs, increasing ocean temperatures are driving large-scale transitions to bleached and low-diversity states dominated by macroalgae (Hughes et al., 2017). Thus, spatial patterning of algae cover in coral reefs may also serve as early warning indicators of catastrophic shifts (Arif et al., 2022; Mumby et al., 2007).

Several mechanisms have been proposed for coral halo formation, where algae cover is reduced, each with a different emphasis on the driving factors. One such mechanism is the landscape of fear (LOF) hypothesis, which suggests that herbivore foraging behavior is shaped by their perceived risk of predation (Gaynor et al., 2019; Laundré et al., 2001; Lima and Dill, 1990). According to this hypothesis, the presence of halos results from herbivores preferentially grazing near the protective cover of coral patches, leading to a clear zone around the coral where algae are sparse.

In contrast to the behavioral focus of the LOF hypothesis, other proposed mechanisms emphasize the role of both endogenous and exogenous factors in spatial pattern formation. For example, reaction-diffusion models can generate both static and dynamic spatial patterns including spots, stripes and spirals that are similar to the skin patterns found in some fish and other animals (Meinhardt, 1999; Turing, 1952). This is thought to arise from the propagation and differential diffusion of pigmented and non-pigmented cells during morphogenesis (Turing, 1952). These reaction-diffusion models rely on endogenous interactions such as those between species (e.g., competition and predation) and their movements to produce spatial patterning (Alevizon, 2002). Nutrient toxicity is one proposed mechanism for coral halo formation, whereby high nutrient loads close to reefs prevent recruitment of algae (Alevizon, 2002). The nutrient toxicity could represent a reaction to the diffusion of algae propagules in a reaction-diffusion context, potentially contributing to spatial pattern formation of halos (Allgeier et al., 2017; Madin et al., 2022). However, exogenous physical factors such as hydrodynamics, substrate characteristics, and light and the availability of other abiotic resources, may also contribute directly or indirectly to halo formation (D’Angelo and Wiedenmann, 2014; Rogers et al., 2014; Steiner and Willette, 2014). For example, Ziemen proposed that halos result from the deposition of larger sediment particles around the coral, which can inhibit algae growth and form a clear zone (Zieman Jr, 1972).

Despite recognizing coral halos as a potential conservation tool, current theories of halo formation have not fully satisfied empirical tests (Bilodeau et al., 2021; Madin et al., 2022). So far, most empirical evidence favors the LOF hypothesis (Madin et al., 2022). Evidence includes documented reductions in algal grazing pressure as distance from coral heads increase on reefs, the ability of coral reefs to provide spatial refugia from predators, and a higher prevalence of halos found on protected reefs in global satellite surveys (Almany, 2004a,b; Madin et al., 2019a). However, some argue that considering consumer-resource interactions, coral halos should be larger where predator populations are reduced, a prediction that is not fully supported by available data. Rather, data indicate that where present in reefs, halos can grow, shrink, and sometimes disappear over space and time but there are no clear links between these halo dynamics, halo widths, and the observed foraging behaviors of herbivores (Madin et al., 2019b, 2022). Halos do appear to have some spatial dependencies: halos are not found consistently across reefs, and in some locations are particularly persistent and prevalent. Halos in the Caribbean were documented in the same location nearly 60 years apart. Halos cover 80% of surveyed reef area in the North Atlantic Ocean, compared to a global average of 10% (Madin et al., 2022). Thus, layering physical constraints on top of consumer-resource and LOF frameworks may help improve predictions of halo widths, persistence, and dynamics.

The spatial distribution of coral patches on a reef is one obvious, yet under-examined physical constraint to halo formation. Coral patches can be highly clustered to dispersed and their distribution is likely to impact both LOF and consumer-resource effects. Thus, we argue that incorporating the effects of static coral distributional patterns are pre-requisite to further understanding the roles of the LOF and consumer-resource interactions in halo formation. Here, we develop a set of simple mathematical models to (1) determine the types of coral spatial patterns that support halo formation using geometric rules and (2) characterize potential interactions between coral distributions and consumer-resource interactions. We then use satellite data from a well-studied reef system for coral halo phenomenon, Heron Island, Australia, as a case study to test whether observed patterns are sufficiently described by geometric rules that assume LOF constraints or whether the inclusion of dynamic consumer-resource interactions is necessary. We show that halo patterns reflect a combination of consumer-resource interactions, LOF behavior, and geometric constraints imposed by static coral reef spatial distributions. These results help to resolve current discrepancies between data and models, underscoring the potential of grazing halos as indicators of ecosystem health and resilience, providing valuable insights for landscape-scale analyses and ecosystem management for reefs and other patchy systems where LOF may apply. We suggest empirical tests that will help to further our understanding of pattern formation under combined LOF, consumer-resource, and habitat clustering mechanisms.

## Methods

We develop two models for coral halo formation: a static geometric model and a dynamic consumer-resource model. From these models, we calculate the predicted long-term fraction of algae for a range of coral spatial distributions. We assume that halos are more detectable as the fraction of algal cover increases across the seascape (Fig. 1). This is because where there is less algae, halos are larger. Where halos are larger, there is more halo overlap, making it more difficult to identify distinctive halo patterns surrounding individual coral clusters. We then compare geometric and consumer-resource model predictions with data on algae and coral distributions derived from satellite imagery of Heron Island, Australia.

### Geometric model: Coral halo geometry and spatial distribution

We consider a square reef of length *L* with a fixed spatial distribution of (identical) coral “units”, interpreted as colonies or fractions of colonies, of radius *r*_*c*_. The spatial distribution of corals ranges from maximally dispersed to maximally clustered, with the average nearest neighbor distance between corals, *δ* ranging from *δ*_*max*_ to 0 (Fig. 2a-b). Though individual corals are of a fixed size, larger clusters represent larger coral colonies or habitat patches for grazing herbivores. Corals are known to provide shelter and protection from predators (Hixon and Beets, 1993). Thus, according to the LOF hypothesis, herbivores will avoid foraging far from coral heads (Ollivier et al., 2018). Though herbivores prefer to graze algae that is closest to coral heads, we assume a well-mixed system where herbivores can move freely across the reef to graze near all coral heads in the reef. Under these assumptions, a grazed algae-free area will form near each coral head: the coral halo.

**Figure 2:**
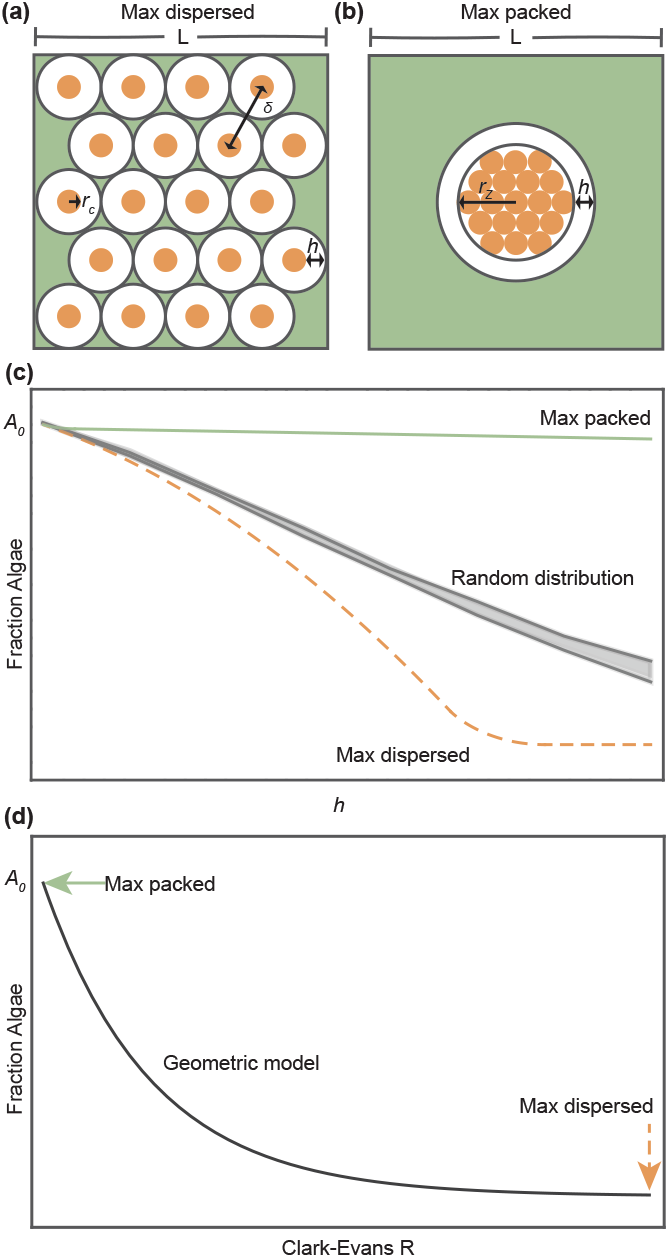
Geometry of coral halos for (a) maximally dispersed (b) and maximally packed coral reefs. Solid orange points represent corals, white annuli are halos. The distance between coral heads is denoted as *δ*, the radius of coral heads is *r*_*c*_, halo width is *h*, and the radius of the circle that encloses the maximum packing of corals is *r*_*z*_. Expected fraction of algae (c) given *h* and (d) the distribution of corals measured in Clark-Evans *R*. In (c), the dashed orange line represents the fraction of algae expected under (a) the scenario where corals are max dispersed and halos do not overlap. The solid green line represents the fraction of algae expected when corals are maximally packed with greatest overlap in halos assuming a coral radius *r*_*c*_ = 1, *n* = 500 corals and a density of corals = 0.0002. Gray band represents the mean fraction of algae +/−2 standard deviations for simulations of randomly distributed corals. Random reefs were generated with n = 500 corals and a square reef of *L* = 1000 pixel width and a density of corals = 0.0002. Random reefs were created for halo widths from h = 0 to 3 in 3/7 intervals with 10 reefs generated for each width. In (d), the solid black line represents the predicted fraction of algae for the geometric model. The dashed orange arrow points to the fraction of algae expected when corals are maximally dispersed and the solid green arrow points to the fraction of algae when corals are maximally packed. In these scenarios, Clark-Evans R is theoretically (a) 2.14 for the maximally dispersed and 0 for the maximally clustered scenarios. *A*_0_ = 1 − fraction of coral, or the fraction of landscape available for algae to colonize. In plot (d), parameter values are *A*_0_ = 0.8, *h* = 3.74, *r*_*c*_ = 2 pixels with *R* ranging from 0-3.

Let us assume that the width of each halo, *h* is fixed to some value, which is dependent on the grazing rate of herbivores in the reef and also on the perceived quality of the coral to act as shelter to herbivores from predators (e.g., different species of coral or fear responses). When two corals are close together, their halos will overlap, reducing the total fraction of free space in the reef generated by the halos. Assuming that algae fills all reef space that is not grazed or occupied by coral, the minimum fraction of reef covered by algae occurs when the corals are maximally clustered (Fig. 2b). When corals are clustered, they form a larger coral head, the area of which can be approximated by encircling an area around the cluster (Fig 2b). Thus, the total fraction of algae for a maximally packed coral reef (*a*_*p*_) is

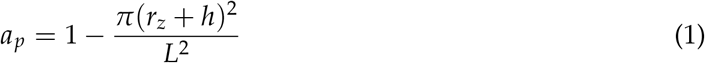

Where *r*_*z*_ is the radius of the encircling area. Conversely, the minimum fraction of algae should occur when the coral are maximally dispersed and *δ* > 2*h*:

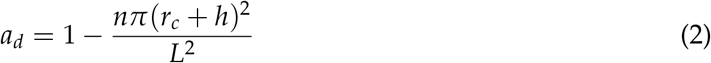

Here, *n* is the number of coral. When *h* = 0, the fraction of algae is *A*_0_, which is the total non-coral fraction in the reef. As *h* increases, the fraction of algae will decline as halos overlap at a rate dependent on the distribution of corals in the reef and the average distance, *δ* between corals (Fig. 2c). We also see that the difference between max dispersed and clustered scenarios increases as *h* increases. To avoid measuring *r*_*z*_, we can use the known density of maximally packed circles to derive a general function that calculates the algae fraction of max packed and dispersed corals as a function of *h* and the density of corals, with details available in Appendix Information A. For simulated random distributions of corals (Fig. 2b), we see that the fraction of algae, *A* at a range of *h* values, can be approximated with an exponential function, where *R* represents the distribution of corals:

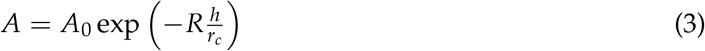

To measure the spatial distribution of corals, we use Clark-Evans R (Clark and Evans, 1954),

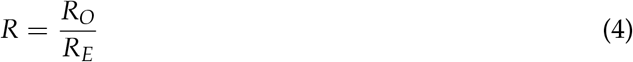

Here, *R*_*O*_ = the observed mean nearest neighbor distance between corals, and *R*_*E*_ = the expected mean nearest neighbor distance for a random Poisson point process of equal intensity. Thus, when the distribution is random, *R* = 1, when it is clustered, *R* < 1, and when it is dispersed *R* > 1.

Our geometric model provides a baseline expectation for how LOF should limit algal growth given the spatial distribution of coral and a set distance limit for foraging grazers. We note that our geometric model predictions apply generally beyond corals to other patchy ecosystems where herbivory is concentrated closest to prey refugia.

### Consumer-resource model

To understand the impact of consumer-resource interactions on halo formation, we develop a spatially implicit consumer-resource model that also incorporates how the LOF and spatial distribution of prey refugia limits growth of primary producers. The geometric model (Eq. 3) predicts the fraction of algae expected when expanding the halo width, *h* given a static spatial distribution of corals, *R*. In this dynamic model, we limit the carrying capacity of algae to this fraction of algae predicted by the geometric model. Since we assume herbivores (e.g., fish) can freely relocate at any coral foci, the dynamics can be described by a mean-field consumer-resource model of algae (A) and herbivores (H) where:

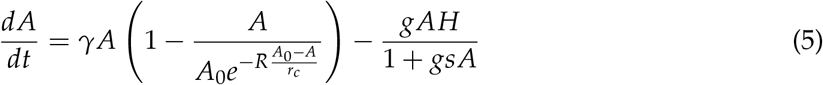

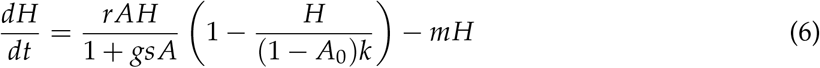

Though the herbivores in our model are mean-field, growth and decline of algae and herbivores are constrained by the spatial distribution of coral, *R*. This approach allows us to include the spatial implications of the LOF and account for the long-distance movement patterns of the herbivores without sacrificing model generality. The model reflects classic consumer-resource assumptions and is broadly applicable to systems where herbivores have long-distance ranges but display some behavioral preferences for grazing near patches of shelter (e.g., elk near forest patches or cows near trees in pastures). We assume that grazing occurs at the boundary of the overlapping circles and that halo width, *h* of Eq. 3 will be some scalar function of the fraction of free space in the reef, (*A*_0_ − *A*) in Eq. 5.

The fraction of algae (*A*) increases by expansion towards the coral at the edge of halos at a rate *γ* and decreases due to herbivores at a rate *g* outside of halos with a type II functional response where *s* is the handling time of herbivores. Herbivore populations grow at a rate *r* and are limited by the carrying capacity, *k*, which is some scalar function of the fraction of corals in the reef (1 − *A*_0_). Herbivores also experience mortality at some rate *m* (Eq. 6).

To analyze the consumer-resource model, we calculated zero growth isoclines for the herbivore population and algae fraction and plotted phase diagrams to determine stability and observe all distinct dynamic behaviors near equilibria (saddle, node, limit cycles, other cycles) possible within realistic bounds for the system (A: 0, *A*_0_) and (*H* ≥ 0) and parameter values (0 < *R* < 2.14, *r*_*c*_, *g, s, k, m, r, γ* > 0). Since our model is deterministic and the number of parameters is large, sensitivity analysis was conducted by varying each parameter separately, while keeping the rest constant to test impacts on system behavior and equilibria (see Appendix C). Equilibria could not be determined analytically (> 3 polynomial), thus were determined numerically as roots of overlapping isoclines. The existence and pattern of bifurcations for numerically determined equilibrium values of *A* were compared as each parameter was changed in a direction that increases herbivore limitation (including increased habitat clustering). Qualitative consensus between bifurcation patterns resulting from similar shifts in isoclines allow us to generalize our results beyond specific parameter value sets to consider herbivore limitation and habitat clustering effects more generally.

### Study area and satellite imagery classification

We focused our data analysis of coral halo patterns on Heron Island, Australia, where halos have been extensively ground-truthed and much information regarding the algae-herbivore-coral dynamics is known (Madin et al., 2011, 2019a, 2022; Ollivier et al., 2018). We downloaded a high resolution image of Heron Island by Maxar Technologies using Google Earth Pro. The image is dated 7/3/2016 (Fig. 1). We first narrowed our analysis to detect halos only in shallow-lagoon habitat using available GIS shapefiles from the Allen Coral Atlas (Kennedy et al., 2021; Lyons et al., 2022). The Allen Coral Atlas is a global map of coral reef habitat, which includes geomorphology that was classified based on Planet Dove 3.7m resolution satellite imagery, wave models, and ecological data. Allen Coral Atlas defines shallow-lagoon habitat as sheltered reef that can be completely or partially enclosed, with a flat bottom and dominated by sediment that is no deeper than 5m (Kennedy et al., 2021; Lyons et al., 2022). Focusing on shallow lagoons allows us to restrict our analysis to areas of similar ecological conditions where halos have been documented and where they are easier to detect given their shallow depth (Madin et al., 2019a, 2022; Ollivier et al., 2018).

All image processing was done in ArcGIS Pro (2.6 − 2.8). We projected the shallow-lagoon polygons to the coordinate system WGS 1984 UTM Zone 56S and georeferenced the Heron Island satellite image, with a resulting raster cell size of 1.67m length and width. With the satellite image clipped to the shallow lagoon habitat, we assigned each raster cell (1.67 x 1.67m) as algae, coral or sand based on an unsupervised classification by spectral signature with manual intervention and additional processing, including accuracy compared to manually classified parcels described in detail in Appendix B.

### Coral spatial distribution from satellite imagery

We sub-sampled the Heron Island reefs by placing square grids at 1X-8X scale, where *X* = 47*m* across the entire shallow lagoon layer. From here, only complete square sub-samples with greater than 0 coral pixels were selected to reduce edge effects and correspond better with model predictions that are formulated for square plots. The spatial distribution of classified coral pixels was measured using Clark-Evans R (Clark and Evans, 1954). We calculated Clark-Evans R for *n* = 1298 at 1X, *n* = 260 at 2X, *n* = 40 at 4X, and *n* = 13 at 8X subplots of classified coral pixels. For each subplot at each spatial sampling scale, we calculated the fraction of pixels classified as coral and algae from the total area sampled. The fraction of coral and algae are equivalent to 1 − *A*_0_ and *A* in our models. This sub-sampling procedure allowed as to account for herbivores that graze at different scales. For example, finer scale spatial features would be more relevant to sea urchins or territorial fish species with high site fidelity while larger scale features would be more relevant to roving fish species (Eynaud et al, 2016).

### Model and spatial data comparisons

We fit our geometric model to spatial data for 1,2,4, and 8X spatial scale sub-samples using maximum likelihood to approximate halo widths. We compared results from 1,2,4, and 8X spatial scales by simulating *n* = 100 reefs using the geometric model and a binomial error distribution with the halo width parameter estimated with maximum likelihood using the quasi-Newton Broyden-Fletcher-Goldfarb-Shanno (BFGS) algorithm for each spatial scale assuming *r*_*c*_ = 2 pixels and starting with an initial halo estimate of 8 pixels. The fit of the geometric model and effects of spatial scale were assessed by drawing means and 95% quartiles of simulated algae fractions for *R* ranging from 0-3 and likelihood ratio tests comparing model fits to null binomial expectations. To compare spatial data results against the consumer-resource model, we tested for multimodality, number of modes, and plotted data assigned into 95% confidence interval mode clusters to assess how significant cluster assignments change with *R* (see Appendix B).

## Results

We found that the geometry of overlapping halos predicts a declining fraction of algae as halo width, *h*, increases and the distribution of corals moves from dispersed to clustered (Fig. 2). This geometric relationship can be approximated with an exponential function for coral spatial distributions between max clustered and dispersed scenarios, with the difference between max clustered and dispersed scenarios increasing with *h* (Eq. 3, Fig. 2c-d).

When split into equally sized square sub-samples, the spatial distribution of corals on Heron Island ranges from *R* = 0.10 to 3.06 (Fig. 3). Increasing dispersion of corals corresponds to a declining fraction of algae as expected given the predictions of the geometric model. When the geometric model is fitted to the data, maximum likelihood estimates of halo width range from 8.44-9.84m across the different sampling spatial scales (Fig. 3c). Simulations of the geometric model with parameterized halo widths assuming binomial error distributions show overlapping 95% quartiles across spatial scales sampled (Fig. 3c). The overlap in confidence intervals indicates that spatial scale does not significantly impact the modeled relationship between coral spatial distribution and algae fraction (Bolker, 2008).

**Figure 3:**
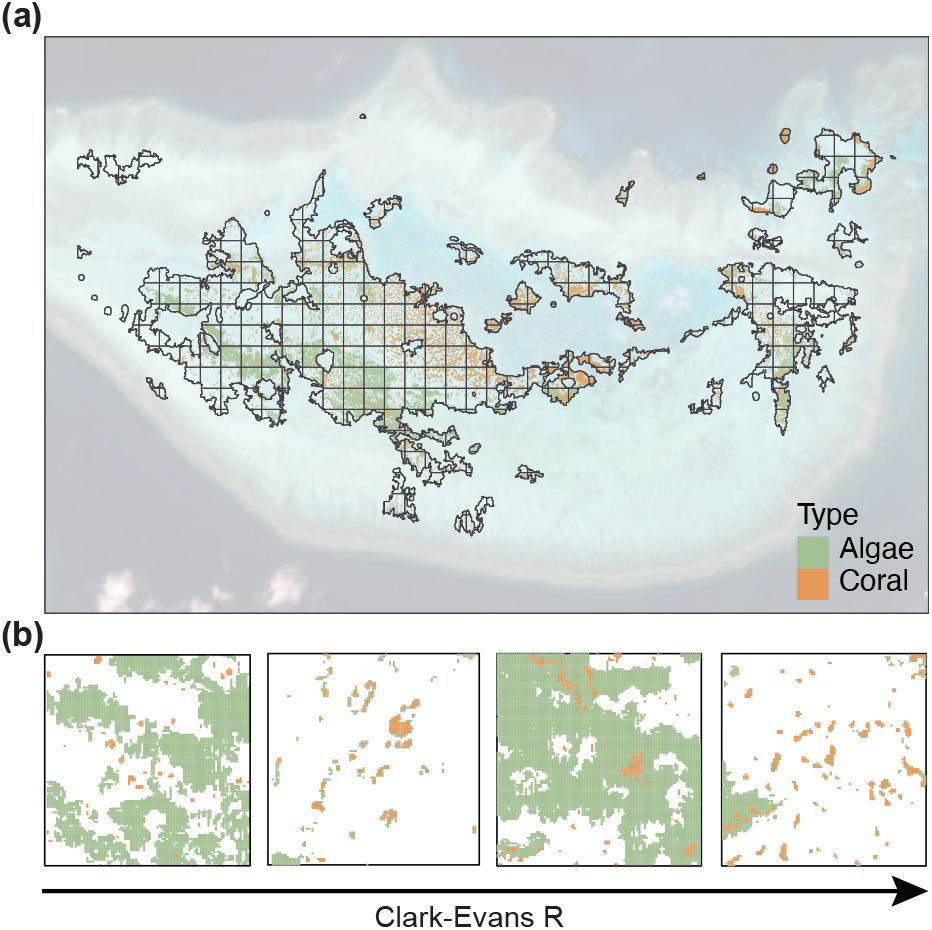
Satellite imagery (background image) of shallow-lagoon layer from Allen Coral Atlas (black boundary) for Heron Island, Australia was classified into (a) algae (green dots) and coral (orange dots) for complete, square sub-samples using a grid of reef drawn at multiple spatial scales. 4X spatial scale of grid is displayed. (b) Example classifications of increasingly dispersed coral (orange dots) reef spatial distributions measured using Clark-Evans *R* (higher values indicate higher dispersion). Reefs sampled at 4X spatial scale. Algae spatial distribution (green dots) also plotted.

Data on actual algae fractions and coral spatial distributions in Heron Island indicate that the geometric model accurately predicts declines in algae as dispersion increases (Fig. 3c, Likelihood ratio test *p* < 0.001 for geometric model at each spatial scale compared to null) (Bolker, 2008). However, especially for more clustered coral reefs, the geometric model fails to capture the spread in data with many algae fraction values falling outside of the 95% confidence bands, indicating that these data do not statistically fit geometric model predictions (Bolker, 2008).

The dynamic consumer-resource model reveals 5 possible equilibria. Two are trivial and unstable (non-varying with parameter value shifts) at (0,0) and (*A*_0_, 0), representing extinction of both herbivores and algae or herbivores alone, respectively (Fig. 4a). Depending on parameter values, three additional equilibria are present. An unstable equilibrium separates a stable node and a cycle, which represent two alternative states: a consistently high algae fraction and a cycle fluctuating close to the origin with periodic declines in algae (Fig. 4a-c). The fraction of free-space can be calculated as *A*_0_ − *A*, which indicate halos that oscillate in size (appearing and reappearing) or stay static over time with a consistently high predicted fraction of algae (Fig. 4d). There is a zone of hysteresis where initial conditions determine whether the algae fraction fluctuates in a cycle or is static at the stable node (Fig. 4d). The system can be forced to move from the cyclic equilibrium where algae fraction fluctuates near zero to a stable node where halos are static for a range of parameter values when *s* (satiation of herbivores), *A*_0_ (fraction space available for algae to colonize), *γ* (growth rate of algae), *m* (mortality of herbivores) are increased or *R* (spatial dispersion of coral), *k* (carrying capacity of herbivores related to coral fraction), *g* (grazing rate of herbivores), and *r* (conversion rate of algae to herbivores) are decreased. These changes all indicate limitations on the herbivore population, a robust and general model result that is independent of changes in specific parameters or parameter values (Fig. 4, see Appendix C).

**Figure 4:**
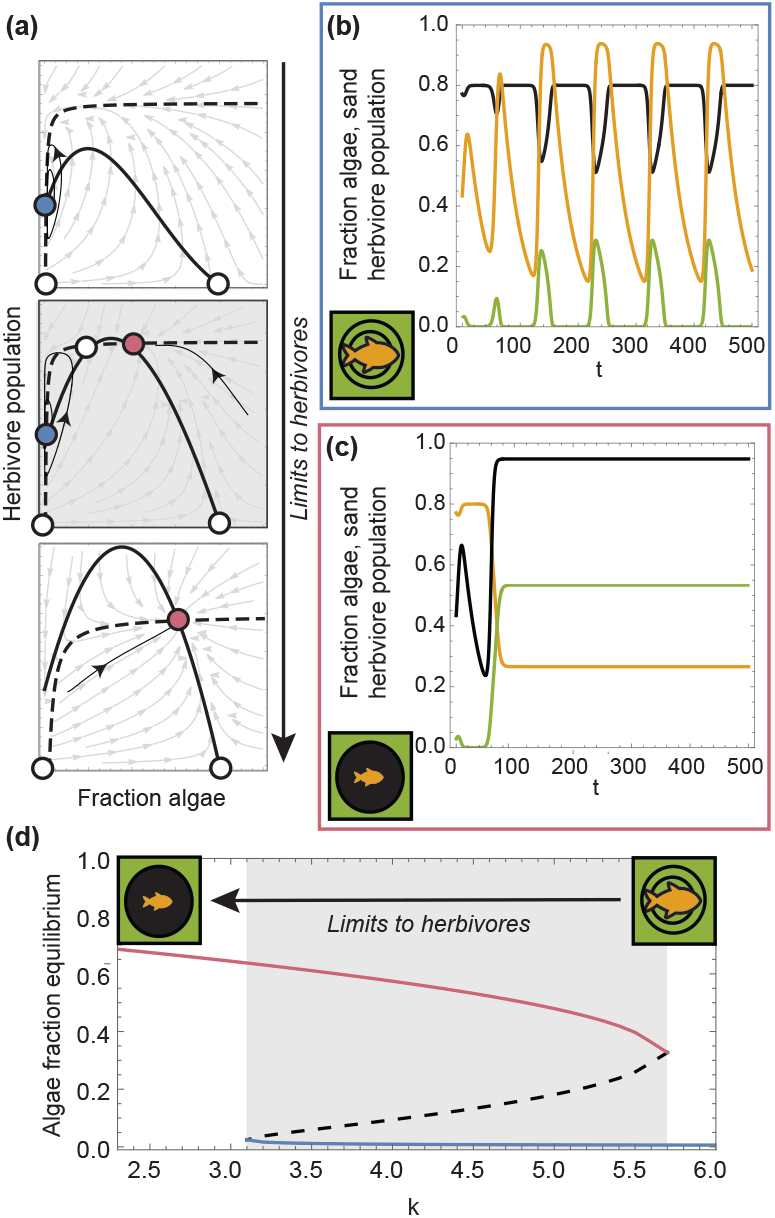
Herbivore model results. Phase diagrams (a) indicating all possible dynamic model behaviors as herbivore (dashed black curve) and algae (solid black curve) isoclines cross producing equilibrium points at their intersections. An additional non-displayed herbivore isocline exists where herbivores are extinct, producing two unstable equilibria at (*A*_0_, 0) and (0,0), represented by open points. Solid blue point represents equilibrium in the center of a cycle with gray vector field indicating direction of flow and solid black arrows representing example trajectories. In gray middle panel, two additional equilibria appear, one open unstable equilibrium point separating a blue equilibrium point with cyclic behavior and an alternative solid pink equilibrium point representing a stable node. Initial conditions determine whether the system moves towards blue or pink equilibria. Increasing herbivore limitation pulls the herbivore isocline down, moving the system from top to bottom panels in (a) between cyclic and stable node behavior. Example time series plots of herbivore population (orange curves) and the fraction of algae (green curves) and free-space (black curves) in (b) cycle where halos oscillate in size or (c) towards stable node with halos moving towards a fixed width. In panel (d), the value of pink, blue, and unstable equilibrium (dashed curve) points are plotted as carrying capacity of herbivores, *k* is varied. Zone of hysteresis is marked by gray box where pink and blue alternative states are present, which corresponds to the gray middle phase diagram panel of (a). In (a), top panel *A*_0_ = 0.8, *R* = 2.14, *g* = 2, *k* = 5, *m* = 0.03, *r* = 8, *s* = 6, *r*_*c*_ = 2, and *γ* = 0.8. For middle panel *R* = 0.92, and for bottom panel, *R* = 0.1 and *m* = 0.14. In (b), parameters are the same as previously except *R* = 1.4, and *m* = 0.03. In (c), *R* = 0.14 and *m* = 0.03. In (d), *R* = 0.5 and *m* = 0.03.

Though we do not have the ecological data necessary to fit the consumer-resource model parameters, patterns in the Heron Island data align with the bifurcation patterns in our consumer-resource model (Fig. 5). The fraction of algae on Heron Island declines exponentially with increasing *R* for all spatial scales sampled (Fig. 4, Likelihood ratio test *p* < 0.001 for 1 − 4*X*). Data sampled at the 1X scale are significantly bimodal (*D* = 0.015, *p* = 0.039) with means (0.043, 0.38) and sd (0.18, 0.080) fraction algae (Fig. 5a) (see Appendix B). When classifying satellite data into significant high and low fraction algae clusters, high fractions occur predominately where coral are more clustered, which is where static halos are predicted by the consumer-resource model. Low and intermediate fractions of algae exist across the range of *R*, as does the cyclic equilibrium in the consumer-resource model (Fig. 5).

**Figure 5:**
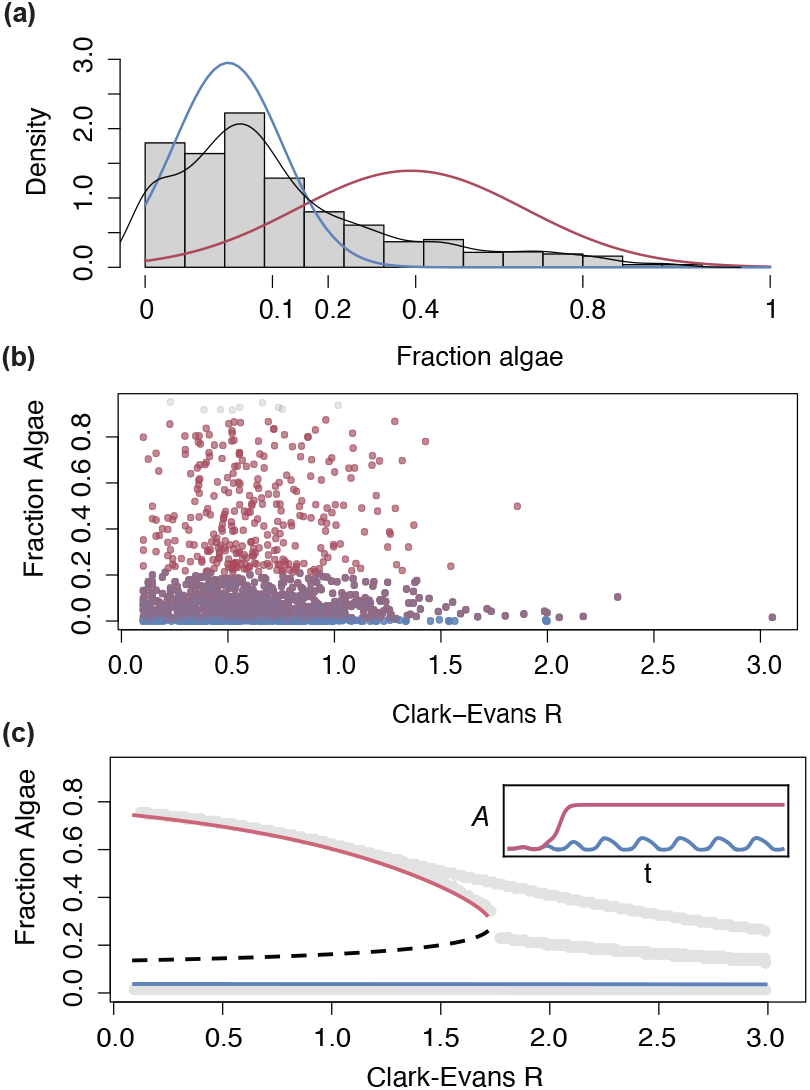
Herbivore model compared to data. Histogram and probability density function (black) of satellite data on fraction algae with fitted Gaussian distributions for two identified modes representing high (pink) and low (blue) fractions. The fraction of algae observed from (b) satellite imagery at 1X scale (47*m*^2^) color coded as belonging in the 95% confidence interval of identified low (blue) and high (pink) fractions, showing intermediate points in purple and outliers in gray. Equilibrium fractions of algae (c) predicted by the consumer-resource model as a function of the spatial distribution of corals measured in Clark-Evans R. Bifurcation occurs as distribution of corals becomes more clustered (R decreases) and a stable node (pink curve) appears as an alternative to cyclic state (blue curve). These alternative states are separated by an unstable equilibrium (dashed curve). Inset demonstrates how the fraction of algae (*A*) changes over time in each alternative state with *A* = fraction of algae. In (c), *A*_0_ = 0.9, *g* = 2, *k* = 4.8, *m* = 0.09, *r* = 8, *s* = 14, *r*_*c*_ = 2, *γ* = 0.15, and R = 0.2 (pink time series) or R = 2 (blue time series) for inset. Gray curves are where *A*_*0*_ (*t*)= 0, which are the peaks and troughs of time series data for the model simulated for 2000 time steps including all transience.

## Discussion

Implementing coral halos as a conservation tool necessitates elucidating the link between coral spatial distributions, consumer-resource reef dynamics, and ecological resilience. We tested two models based on the LOF hypothesis: a geometric model that implicitly assumes LOF and a dynamic mean-field model with explicit consumer-resource dynamics. The geometric model predicts empirically observed decreasing macroalgae fractions when coral patches are more dispersed (higher values of *R*). The dynamic model with macroalgae-herbivore dynamics further explains observed patterns, namely high variability of the macroalgae fraction when coral have high and intermediate clustering (low to intermediate values of *R*). When coral are clustered, the system can exhibit a high and stable algal cover, oscillations around low algal cover, and bistability between these two states (Figs. 5). Overall, we find that the utility of halo imagery as an indicator of ecological resilience is highly dependent on coral spatial clustering and the parameters associated with characteristics of the algal and herbivore communities.

We found that geometric rules can explain the lack of halos in dispersed coral reefs, which is also consistent with LOF theory (Lima and Dill, 1990). Although this model only implicitly assumes LOF, it accurately predicts halo widths corresponding to previously published results on Heron Island (Madin et al., 2019b; Ollivier et al., 2018). We note that halo width may be a useful measure of habitat patch quality or fear (e.g., different coral species, health status, etc.) but only where halos are temporally persistent. Satellite imagery rastered to a resolution of 1.67m (currently freely available through Google Earth) was an effective tool for monitoring halo formation and size, at least for Heron Island.

Modeling consumer-resource interactions helps to explain patterns in the satellite data that deviate from geometric model predictions. The data is bimodal, supporting the existence of alternative stable states predicted by our consumer-resource model. Furthermore, the overarching patterns in the data showing low and intermediate fractions of algae across the range of *R*, and high fractions limited to parcels with the most clustered corals, are consistent with our modeled bifurcation pattern. In these results, two alternative states (stable halos and cycles) collapse into one state (cycles) as *R* increases and herbivore limitation declines. The observed intermediate algae fractions from satellite data, which are not significantly different from high and low clusters, are consistent with dynamic expectations for points sampled in transition between alternative states for several reasons. First, dynamics are necessarily slower near both unstable and stable equilibria (Strogatz, 2018). Second, stochasticity can lead to flips between alternative regimes and we detected two statistically significant modes, not three (see Appendix B). Third, the low algae equilibrium is cyclic and we do not know how far from equilibrium the system is at the point of sampling. In Fig. 5c, we show the peaks and troughs of time series data, which illustrates that the stable node can act as a “ghost” attractor even past the critical *R* value where it is eliminated due to transient dynamics, which may account for some points being classified as intermediate and high algae fractions when coral are more highly dispersed.

Previous work on coral reef alternative stable states shows that the grazing intensity of herbivores and initial conditions determine whether the system will reach a high coral/low algae state vs. a low coral/high algae state (Mumby et al., 2007). Our results provide evidence that LOF can also lead to alternative regimes in coral reefs, helping to explain the heterogeneity of algal cover observed over time and across reef systems (Madin et al., 2019a). Because our framework assumes that corals are static, our findings best represent reef systems where algae grow much faster than coral while the Mumby et al. (2007) model best approximates systems where coral and algal growth rates are more similar. We note the importance of these timescale considerations: under similar coral and algal growth rates, high spatial clustering can decrease the potential for bistability through increased intraspecific competition (Tekwa et al., 2021) while we find that bistability only occurs when our static corals are clustered.

Assuming that halos are more detectable at higher algal cover, we find that stable halos, halos which increase and decrease cyclically over time, and the absence of halos are all consistent with LOF theory. These results may help contextualize previous findings that linked halo size to patch reef area but not to marine reserve status in the Great Barrier Reef (Madin et al., 2019b). The missing link between halo size and reserve status is usually cited as evidence against LOF theory but our model provides an alternative explanation, namely, that larger patch reefs could indicate more clustered coral colonies that increase the potential for high algal cover. Future research that controls for coral spatial distribution (e.g., looking at similarly clustered reefs) while varying marine reserve status would help refine tests of the LOF hypothesis. Additionally, our consumer-resource model predicts that increasing limitations to herbivores (e.g., in protected reefs with predators such as sharks or under climate change scenarios) can drive the system towards a stable node where halos are more persistent and easier to detect. This is consistent with research that linked marine reserve status to the age of halos (older halos were found in reserves) rather than their size (Madin et al., 2019b). This same research detected halo “footprints”, potentially indicating long-term cycling behavior that is one alternative state in our model. Though we predict more abundant predator populations to induce more persistent halos, long-term dynamics are also important to consider. If herbivore populations are overexploited, predator extinction can occur, resulting in algae dominance with zero free space (i.e., no halos, only algae and corals) (Fig. 4). (Hughes et al., 2003). However, our model does not yet incorporate dynamic predator populations or consider metapopulation dynamics of more constrained herbivore populations, areas we suggest for future research.

Empirical studies focused on the predominance, amplitude, and frequency of cyclic behavior in halos may provide further insights into the health of consumer-resource interactions. Where cyclic behavior is present, we expect that tracking the amplitude and frequency of cycles would be useful for assaying the strength of herbivore limitation, though transitions to alternative stable node dynamics may complicate correlative analyses (Fig. 4-5). We find that limitations on the herbivore population drive shifts from cyclic to static halos. Thus, increased prevalence of static halos is expected for protected reefs. The scale of LOF (i.e., how much risk is perceived by herbivores) may also influence herbivore limitation, with more risk expected to result in more consistent halos by reducing grazing rates.

Although we assess our models with data from Heron Island, they are constructed to be broadly applicable to any ecological system where LOF dynamics and patchy habitats are relevant. We find that static spatial pattern formation in primary producers resulting from LOF is only expected where protective patches are relatively clustered. Frequency of grazing halos on reefs and other patchy systems is still a useful indicator of LOF-induced herbivore limitation but can only be correctly compared across sites where refugia are sufficiently and similarly clustered. For example, static fairy circles produced by termites are predicted to be present only where nests are clustered and if nests protect from predators. These predictions provide clear testable hypotheses for future research. Since both static and dynamic pattern formation are consistent with LOF theory, failure to detect static pattern formation cannot signal low herbivore limitation or exclude the applicability of LOF processes. Thus, these processes may be more broadly relevant than previously realized, especially where the detection of static spatial patterns was used as primary evidence. Still, the existence of temporally persistent grazing halos, whether on reefs or elsewhere, does signal strong herbivore limitation and the presence of LOF. Additionally, we find that herbivore limitation in such systems can drive alternative stable state behavior relevant to ecosystems with managed herbivore populations. A sudden loss of formerly persistent grazing halos provides an important signal of a critical transition in herbivores that will be difficult to manage for recovery. Persistent grazing halos may be a key characteristic used to identify sites for long-term monitoring of ecological resilience.

## Conclusion

As a reef manager, maintaining more persistent and consistent herbivore populations and halos could be desirable (assuming that the clustering of coral patches supports halo formation) (Mumby and Steneck, 2008; Mumby et al., 2014). Future research could assess whether reefs that have more persistent halos also have more stable herbivore and predator populations. Additionally, experimental changes such as stocking algae or herbivores could test whether stable halos can be induced in cyclic systems as part of management strategies (Hughes et al., 2003, 2017). Taken together, this study and the future directions we highlight represent promising advances towards linking satellite imagery, halo patterns, and reef community dynamics to support reef conservation efforts and have broad relevance for pattern formation in other patchy ecosystems limited by LOF.

## Acknowledgments

We thank Alana Danieu for help with preliminary analyses, Ashley Laveriano for manually classifying pixels, and the Elizabeth Madin Lab Group for helpful feedback and discussions.

## Statement of Authorship

TWO, LM, VVV, and LY jointly designed the study, performed the research, developed and analyzed models, produced figures, and wrote the manuscript. CS processed satellite imagery data. TWO analyzed spatial data and led first draft of manuscript. All authors contributed substantially to revisions.

## Data and Code Availability

The data supporting the results and the code used to generate the figures are archived in Zenodo, which will be released for open access following acceptance of manuscript. The files are available for review at the following DOI: https://zenodo.org/doi/10.5281/zenodo.8335656.

