## Appendices for "Seeing halos: Spatial and consumer-resource constraints to landscape of fear patterns"

### Appendix A: Coral halo geometries

In this appendix, we provide the formulas that bound the fractional algae density for a fixed number of coral heads, $n$, coral radius, $r_c$, and halo width, $h$. As illustrated in Fig. 2, those correspond to the maximally and minimally dispersed cases for the coral heads.

#### *Maximally dispersed coral heads*

Assume all corals are the same size, $r_c$, and equally (and maximally) spaced, configured in a hexagonal lattice, in a square reef of area $L^2$ (Fig. 2a-b). Let $\delta$ be the distance between the center of two consecutive corals. Let $r_t = r_c + h = \alpha r_c$ be the radius where fish can graze (including coral radius, $r_c$, imposing $\alpha \geq 1$) and where the amount of algae is negligible. We call $r_t$ the radius of the halo and $h$ the halo width. When the radius of the halos, $r_t$, is less than half of the distance between two consecutive corals, $\delta/2$, there is no overlap between halos and algae grows outside $n$ isolated circles, each of area $\pi r_t^2$. Above that, the halos overlap. When the halos touch the center of the triangles formed by the coral heads, there is no more algae left. The center of the triangle is at a distance $\frac{2\sqrt{3}}{6}\delta$ from the center of the corals (the vertices of the triangle), so there is no algae for $r_t$ larger than that. In between, i.e., for $\frac{\delta}{2} < r_t < \frac{2\sqrt{3}}{6}\delta$, each halo overlaps equally with 6 other halos of the nearest coral heads. To compute the area of algae, we need to add the overlap between those circles to the subtraction of the area of the $n$ circles. The area of two overlapping circles is $2\left(r_t^2 \cos^{-1}\frac{\delta}{2r_t} - \frac{\delta}{4}\sqrt{4r_t^2 - \delta^2}\right)$.

Thus, the fraction of area covered by algae is given by

$$a[n, r_t, L, \delta] = \begin{cases} 1 - \frac{n\pi r_t^2}{L^2}, & r_t \leq \frac{\delta}{2} \\ 1 - \frac{n\pi r_t^2}{L^2} + \frac{6n}{L^2}\left(r_t^2 \cos^{-1}\frac{\delta}{2r_t} - \frac{\delta}{4}\sqrt{4r_t^2 - \delta^2}\right), & \frac{\delta}{2} < r_t < \frac{2\sqrt{3}}{6}\delta \\ 0, & r_t \geq \frac{2\sqrt{3}}{6}\delta \end{cases} \tag{A1}$$

Now, the number of coral heads, $n$, in a square area of size $L^2$ and the distance between coral heads, $\delta$, packed in a hexagonal lattice are not independent. Let us denote the density of coral

$f = n\rho = n\frac{\pi r_c^2}{L^2}$ and the relative spacing between corals $\Delta = \frac{\delta}{2r_c}$. Then, the fractional area of algae in the gaps of hexagonally packed corals can be rewritten as

$$a[f,\Delta,\alpha] = \begin{cases} 0, & \frac{\Delta}{\alpha} \leq \frac{\sqrt{3}}{2} \\ 1-\alpha^2 f + \frac{6\alpha^2 f}{\pi}(\cos^{-1}\frac{\Delta}{\alpha} - \frac{\Delta}{\alpha}\sqrt{1-(\frac{\Delta}{\alpha})^2}), & \frac{\sqrt{3}}{2} < \frac{\Delta}{\alpha} < 1 \\ 1-\alpha^2 f, & \frac{\Delta}{\alpha} \geq 1 \end{cases} \tag{A2}$$

Finally, since the minimal fraction of algae occurs when corals are hexagonally packed in the entire space (Fig. 2a), and the density of that packing is

$$\eta = \frac{n\pi\frac{\delta}{2}^2}{L^2} = \frac{\pi\sqrt{3}}{6}, \tag{A3}$$

the distance between coral foci under hexagonal packing (maximum density) is

$$\delta = \frac{L}{\sqrt{n}}\frac{\sqrt{2}}{3^{\frac{1}{4}}} = c\frac{L}{\sqrt{n}}, \text{with } c = \frac{\sqrt{2}}{3^{\frac{1}{4}}} \approx 1.07. \tag{A4}$$

So, $\Delta = \frac{c}{2}\sqrt{\frac{\pi}{f}}$, and the minimum fraction of algae we should see in a reef

$$a_{min} = a\left[f, \frac{c}{2}\sqrt{\frac{\pi}{f}}, \alpha\right]. \tag{A5}$$

#### *Minimally dispersed coral heads*

The maximum fraction of algae corresponds to when the corals are clustered together (given fixed number of coral, $n$, and fixed reef area, $L^2$), or $\delta = 2r_c$. Assume the corals cluster hexagonally in a large circle, $Z$, of radius $r_z$, made of the $n$ coral heads with area $C$. Then, the fraction of space covered by algae in the entire square space is (Fig. 2b)

$$a_{max} = 1 - \frac{C}{L^2} + (\text{fraction of algae in } Z)\frac{C}{L^2} - \frac{\text{area of grazing halo surrounding } Z}{L^2}. \tag{A6}$$

The radius of the coral circle $Z$, $r_z$ can be calculated as a function of the number of coral heads and their individual radius from the density of hexagonal packing

$$\eta = \frac{\pi\sqrt{3}}{6} = \frac{n\pi r_c^2}{\pi r_z^2}. \tag{A7}$$

Therefore,

$$\begin{aligned}\frac{C}{L^2} &= \frac{\pi r_z^2}{L^2} \\ &= \frac{2\sqrt{3} n r_c^2}{L^2} \\ &= \frac{2\sqrt{3} n \rho}{\pi}.\end{aligned} \tag{A8}$$

The area of the grazing halo surrounding $Z$ is

$$\begin{aligned}&\pi \left( (r_z + h)^2 - r_z^2 \right) \\ &= \pi \left( (r_z + (\alpha - 1) r_c)^2 - r_z^2 \right) \\ &= \pi \left( 2(\alpha - 1) r_z r_c + (\alpha - 1)^2 r_c^2 \right) \\ &= \pi \left( 2(\alpha - 1) \sqrt{\frac{n}{\eta}} r_c^2 + (\alpha - 1)^2 r_c^2 \right) \\ &= (\alpha - 1) \pi r_c^2 \left( 2\sqrt{\frac{n}{\eta}} + (\alpha - 1) \right).\end{aligned} \tag{A9}$$

So, the maximum fraction of algae cover is

$$\begin{aligned}a_{max} &= 1 - \frac{2\sqrt{3} n \rho}{\pi} + \frac{2\sqrt{3} n \rho}{\pi} a \left[ \frac{\pi\sqrt{3}}{6}, 1, \alpha \right] - (\alpha - 1) \frac{\pi r_c^2}{L^2} \left( 2\sqrt{\frac{n}{\eta}} + (\alpha - 1) \right) \\ &= 1 - \frac{2\sqrt{3} n \rho}{\pi} + \frac{2\sqrt{3} n \rho}{\pi} a \left[ \frac{\pi\sqrt{3}}{6}, 1, \alpha \right] - (\alpha - 1) \rho \left( 2\sqrt{\frac{n}{\eta}} + (\alpha - 1) \right)\end{aligned} \tag{A10}$$

### Appendix B: Satellite imagery analysis

In this appendix, we provide detailed satellite imagery analysis methods including classification of imagery into benthic types, spatial correlation between coral and algae, and analysis of multimodality.

#### *Classification of imagery into benthic types*

We first ran unsupervised classification (Iso cluster) of the shallow lagoon layer to classify raster cells into different classes. To capture fine spectral differences of coral, algae and sand, we specified 10 classes for detecting various spectral signatures, with 20 as the minimum number of

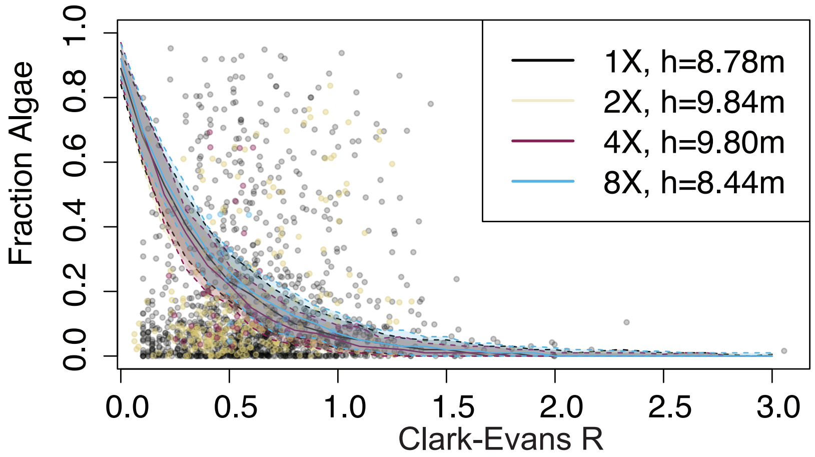


Figure B1: (Relationship between fraction of algae and Clark-Evans $R$ for data extracted at 1,2,4, and 8X spatial scales where $X = 47m$. Solid curves represent medians for $n = 100$ simulations of geometric model with a binomial error. Dashed lines and confidence intervals represent 95% quartiles of model simulations, indicating poor fit with data. Halo widths, $h$ used in geometric model simulations were parameterized at each spatial scale using maximum likelihood fits to data assuming $r_c = 2$ pixels, giving similar halo widths corresponding to emprirical measurements at all spatial scales assessed.

cells in a valid class and a sample interval of 10. From the initial result, we noticed inaccurate and mixed classifications in the lower half of the imagery between corals and algae, due to their similar spectral ranges when compared to the upper half of the image where corals have a more distinctive spectral signature due to potential differences in the sediment ratio. Thus, we split the image (Fig. B1) by preserving the results from the upper half of the image, and extracting the mix-classifications of algae and coral in the lower half of the image. We ran segment mean shift on the mixed classification of coral and algae to group adjacent pixels with similar spectral characteristics for better identification of objects. We then re-ran the unsupervised classification on the extracted layer with the segmented mean shift. We combined the two sections of the image and coded the ten classes to three for algae, coral and sand. The final classifications with spatial information (x, y coordinates) were exported for spatial clustering analysis.

We randomly selected a 28 by 29 pixel rectangular region in the upper and lower half to manually classify. We classified pixels into coral and algae cells, presuming all other cells are sand. Unsupervised classification (upper half) was 92% accurate for corals, 50% for algae, and 41% for sand on upper half without segment mean shift and 87% accurate for corals, 94% for algae, and 77% for sand in lower half plot with shift (Mcnemar's Test $p < 0.05$ for each confusion matrix). To test ambiguity in manual classifications, we asked a person with no expert knowledge of the coral system to indicate black cells as coral, green cells as algae, and presume all other cells are blue sand. Non-expert classifications were 98%, 94%, and 94% accurate for coral, algae, and sand in upper plot and 99%, 100%, and 100% accurate in lower plot when assuming expert classifications (lead author) are true ($p < 0.05$) indicating low ambiguity for observers. These accuracy values are within the range of published classifications of coral halo patterns including more complex deep learning classifications (Franceschini et al., 2023).

#### *Spatial correlation between coral and algae*

To statistically assay halo phenomenon, we ran Cross K-function analysis, which is an extension of Ripley's K for a bivariate Poisson point process to determine how coral and algae are

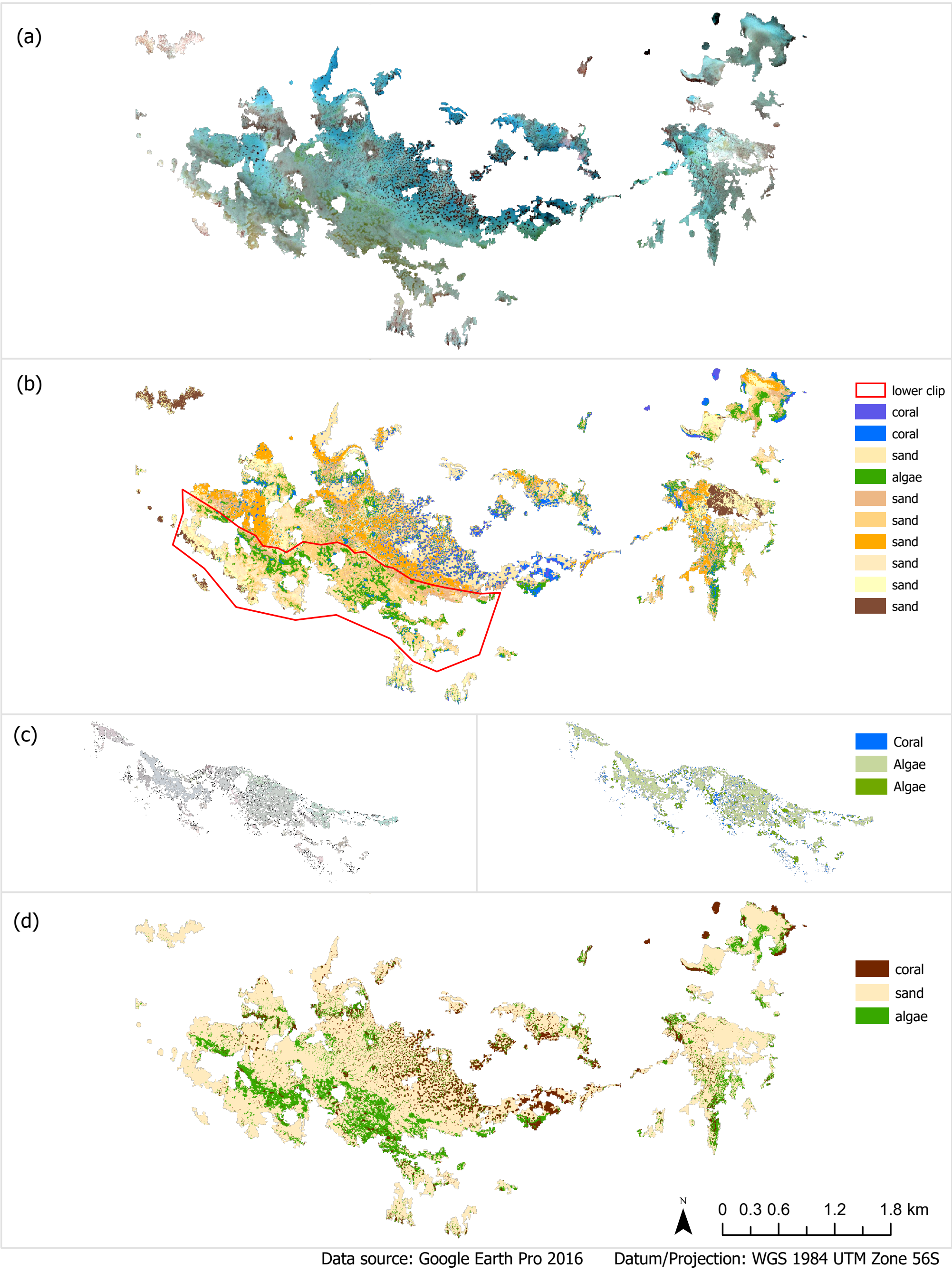


Figure B2: Satellite image classification for Heron Island. (a) Clipped satellite image to shallow lagoon in Heron Island. (b) Initial unsupervised classification with 10 classes of the satellite imagery. Red box marks the area extracted for further classification in coral and algae mixed class. (c) Left image shows results from segmented mean shift on extracted coral and algae mixed class in the lower half of the image, and right image shows reclassification of the area incorporating the segmented mean shift. (d) Final combined image of the classifications to three classes: sand, coral, and algae. The results were exported with geographical information (cell location) and classes (cell identity).

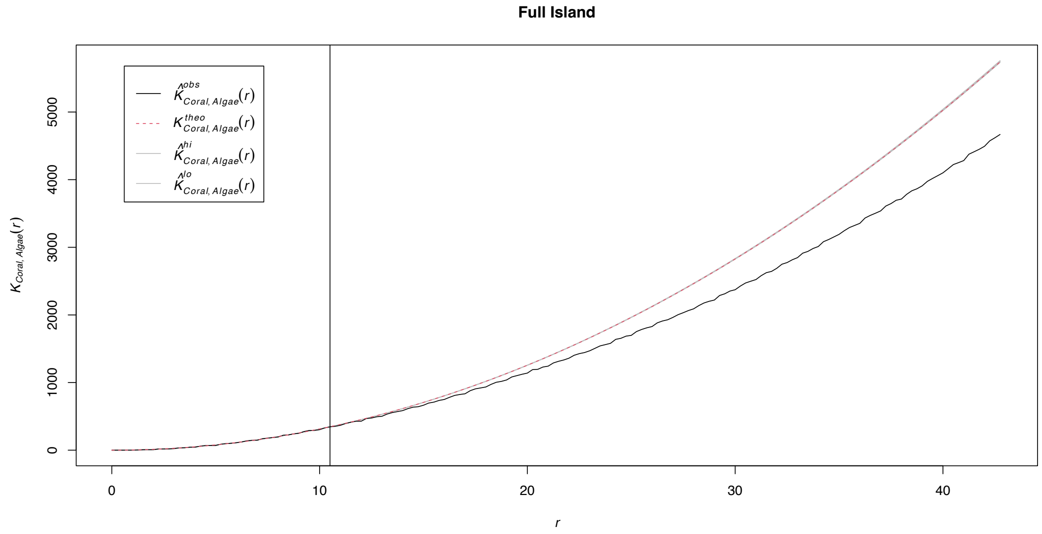


Figure B3: Cross K-function analysis. Beyond a certain spatial scale marked by vertical line, algae are are detected to be more dispersed from coral than expected under a random Poisson process (red curve with gray 95% confidence band), indicating that coral and macroalgae are "repelled" from each other as is the expectation if coral halos are present. Note that this analysis is conducted with data across the entire Heron Island, using a border correction for the shallow lagoon polygon. When considering Heron Island as a whole, there is significant evidence of halos, as also confirmed by other studies where Heron Island is used as a model location for halo phenomenon.

distributed relative to one another (Fig. B2).

#### *Multimodality analysis*

We ran Hartingan's dip tests that confirm multimodality of the data from sampling at the X1 spatial scale where we had most points/power (D = 0.015035, p-value = 0.04695). Following (Hirota et al., 2011), we ran latent class analysis using the FlexMix R package, which conducts expectation-maximization (EM) to assess modality in the univariate normal distribution of algae fraction after an arcsin transformation as a test for the existence and number of alternative stable states. The model with 3 modes failed to converge after n=1000 repetitions, so we raised the minimum prior probability of clusters to 0.2. Components falling below this threshold are removed during the iteration. We compared models for 1-3 classes using Akaike Information Criterion (AIC), Bayesian Information Criterion (BIC), and the Integrated Completed Likelihood (ICL) (Table S1).

Table S1: Goodness-of-fit for models of algae fraction with 1-3 latent classes. The minimal value (bold) indicates best fit for Akaike Information Criterion (AIC), Bayesian Information Criterion (BIC), and the Integrated Completed Likelihood (ICL) within 2 point differences.

| **Criterion** | **1 Class** | **2 Classes** | **3 Classes** |
|---|---|---|---|
| AIC | 413.05 | **423.39** | 423.39 |
| BIC | 423.39 | **-3.64** | -3.64 |
| ICL | 423.39 | **268.27** | 268.28 |

The 2 and 3-classes model were not more than a 2 point difference apart for all criteria tested, the generally accepted threshold for improved fit (Table S1). Both 2 and 3-classes were significantly better than 1 class. Since the 3-classes model was difficult to converge, is more complex, and did not provide significant change in criteria, we considered the simpler 2-classes model the best fit overall. Fitted parameters for the two modes had means= 0.67, 0.21, and standard deviations=0.29, and 0.14 for fraction algae (transformed values). Fig. 5a shows histogram, probability density, and the two fitted normal distributions for fraction algae after arcsin transformation (note the scale of x axis). The detection of 2 modes in the satellite data of algae fraction aligns with the existence of alternative stable states predicted by our consumer-resource model. Since the lower alternative state is cyclic, high variance in single time snapshot data is expected. Corresponding 95% confidence interval clusters for the two modes are plotted in Fig. 5 as a function of $R$ to further statistically support match between the data and the qualitative bifurcation pattern predicted by the consumer-resource model.

### Appendix C: Sensitivity analysis for consumer-resource model

The following appendix provides full sensitivity analysis for consumer-resource model showing the range of bifurcation patterns possible when tuning each parameter separately while controlling all others (Fig. C1)

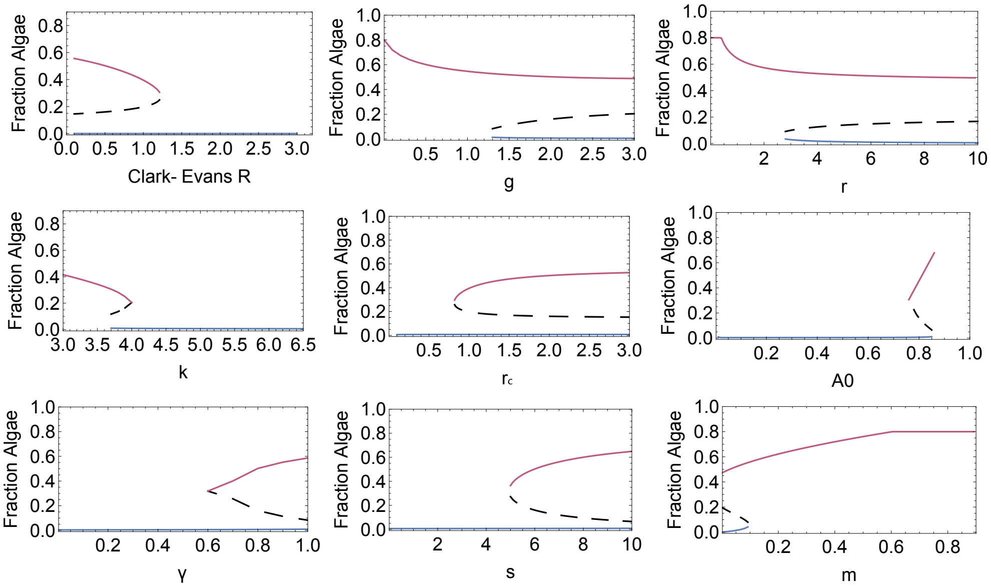


Figure C1: Resulting bifurcations when varying each parameter in consumer-resource model, following the one-at-a-time approach. Parameters (other than what is changing): $A_0 = 0.8, g = 2, k = 4.8, m = 0.03, r = 8, s = 6, r_c = 2, \gamma = 0.8$, $R = 0.5$. $A_0$ is fraction of non-coral space, $g$ is grazing rate of herbivores, $k$ is carrying capacity of herbivores, $m$ is the herbivore mortality rate, $r$ is growth rate of herbivores, $s$ is satiation rate of herbivores, $r_c$ is the radius of individual coral, $\gamma$ is the growth rate of algae, and Clark-Evans $R$ measures the spatial distribution of coral where higher $R$ are more dispersed. Black dashed curve is unstable equilibrium. Pink curve is stable node. Blue curve is cyclic equilibrium.
